# Degeneracy in neuronal bifurcation landscapes: equivalent bifurcation sequences across distinct parameter sets

**DOI:** 10.1101/2025.08.04.668451

**Authors:** Gianmarco Cafaro, Christophe Bernard, Meysam Hashemi, Damien Depannemaecker

## Abstract

Degeneracy—the capacity of structurally distinct systems to achieve similar functions—is a fundamental property of living organisms, enabling adaptability and resilience. Neurons can maintain stable activity patterns despite wide variability in ion channel expression, highlighting how distinct internal configurations can yield equivalent electrophysiological behaviors. However, it remains unclear how degeneracy manifests in the context of bifurcations, the critical transitions between activity regimes that underlie phenomena such as different firing patterns, seizures and depolarization block. Here, we investigate how different biophysical parameter sets can lead to equivalent bifurcation sequences. Using a minimal neuron model with three variable conductances, we systematically explore how changes in extracellular potassium—mimicking physiological and pathological conditions—affect neuronal dynamics. Our results reveal that neurons with distinct intrinsic properties can traverse the same bifurcation pathways, entering regimes of bursting, seizure-like activity, and depolarization block. Yet, the specific parameter set determines the sensitivity and thresholds for these transitions. This work clarifies how degeneracy extends to the dynamical landscape of neurons, with implications for understanding resilience and vulnerability in neural circuits.

**Author Summary:** Biophysical models expressed through differential equations can reproduce the dynamics of biological systems with varying levels of detail. By changing model parameters, simulations can capture the natural variability that allows biological systems to achieve the same behavior through different mechanisms. This property, called degeneracy, underlies the robustness of living systems to internal and external perturbations. In this work, we identify dis-tinct behaviors in single-neuron models and link them to electrophysiological patterns observed under elevated extracellular potassium during epileptic events. These patterns correspond to specific classes of spontaneous bursting activity within the framework of nonlinear dynamics. We show that different combinations of conductance parameters, at a fixed extracellular potassium level, can generate the same class of patterns, defining degeneracy groups. Finally, we assess the robustness of these groups by analyzing how they respond to changes in extracellular potassium. Our findings provide a basis for studying variability and resilience in the dynamics of more complex neuronal systems.

## 2 Introduction

Degeneracy—the ability of structurally distinct systems to produce similar functional outcomes—is a pervasive feature of biological systems [9]. In the nervous system, this property enables neurons to maintain robust activity despite substantial variability in ion channel expression, synaptic connectivity, and membrane properties [1, 22]. Seminal studies by Eve Marder and colleagues demonstrated that individual neurons can compensate for perturbations in internal parameters—such as ionic conductances or channel densities—while preserving stable firing patterns [21, 28]. This flexibility is thought to underlie the resilience of neural circuits to genetic, developmental, and environmental variability. Likewise, the initial configuration of a neuron’s parameters determines how it will respond to external modulations such as temperature, pH, or ionic concentrations [2].

While degeneracy in steady-state or repetitive spiking activity is well characterized, far less is known about how it manifests in the transitions between activity regimes—so-called bifurcations—that are central to both normal and pathological dynamics. In epilepsy and related disorders, neurons often display abrupt shifts in excitability, transitioning between silence, bursting, seizure-like discharges, and depolarization block [18, 32]. These transitions can be mathematically described as bifurcations in nonlinear dynamical systems, where changes in a control parameter (such as extracellular potassium) lead to qualitative changes in neuronal behavior [7, 17].

Epileptic activity has been observed across a wide range of species and brain structures [16, 24, 29], and may arise from diverse etiologies including genetic, traumatic, or metabolic origins [3, 12, 30]. Despite this heterogeneity, electrophysiological signatures such as rhythmic bursting or sustained ictal activity are remarkably conserved. Canonical dynamical models reproduce these activity patterns using low-dimensional systems of differential equations that capture the onset and offset bifurcations defining epileptiform transitions [18, 31]. These phenomenological models can be linked to biophysically grounded descriptions, which incorporate specific ionic currents and concentration dynamics [5–7].

A central question is whether different biophysical implementations—distinct combinations of ionic conductances—can give rise to the same sequence of bifurcations under changing environmental conditions. In particular, how does degeneracy manifest when neurons are exposed to elevated extracellular potassium concentration, a hallmark of intense neuronal activity and epileptic seizures [10, 13, 33, 34]?

To address this, we employ a simplified single-compartment neuron model with three tunable conductances (sodium, potassium, and chloride). By systematically varying both the intrinsic parameters and the extracellular potassium level, we map the resulting activity patterns and bifurcation sequences. Our approach allows us to identify parameter sets that yield equivalent dynamical transitions—e.g., from tonic spiking to bursting or depolarization block—determine how sensitivity to external perturbations depends on intrinsic configuration, and characterize how degeneracy evolves under changing environmental constraints.

This analysis provides insight into the interplay between internal variability and external modulation in shaping neuronal dynamics. While degenerate configurations may exhibit equivalent behavior under baseline conditions, they may diverge in their responses to perturbations—an effect critical for understanding differential treatment responses and drug resistance in epilepsy [27]. By clarifying how different parameter sets produce similar or divergent transitions, our findings help elucidate mechanisms underlying pathological activity patterns and suggest strategies for controlling neural excitability based on intrinsic neuronal state.

## 3 Results

### 3.1 Similar Dynamics Across Parametrization

Abnormal patterns of brain activity are a hallmark of several neurological and psychiatric conditions, including epilepsy (Perucca et al., 2014 [26]), Parkinson’s disease (Levy et al., 2000 [20]), and schizophrenia (Uhlhaas et al., 2009 [35]). A common electrophysiological feature across these disorders is bursting activity, which consists of alternating periods of rapid neuronal firing and quiescence (Izhikevich, 2007 [17]; Saggio et al., 2017 [32]). These bursts—defined as clusters of spikes separated by silent phases—can be categorized into distinct patterns based on properties such as spike amplitude, frequency, and burst duration.

In epilepsy, seizures corresponds to specific types of bursting patterns and are identified in both experimental recordings and theoretical models (Jirsa et al., 2014 [18]; Saggio et al., 2020 [31]). These patterns have been observed across preparations, including in vivo and in vitro studies, and across species, suggesting a conserved underlying mechanism. Importantly, such activity can be understood within the framework of nonlinear dynamical systems, where the transitions into and out of bursting regimes are governed by bifurcations—qualitative changes in the system’s stability structure (Izhikevich, 2007 [17]).

In low-dimensional models, especially two-dimensional systems, the majority of bursting patterns relevant to epileptiform activity are associated with specific combinations of onset and offset bifurcations (Depannemaecker et al., 2022 [7]; Jirsa et al., 2014 [18]; Saggio et al., 2020 [31]). These bifurcations are driven by variations in system parameters, such as ionic conductances, that control the transitions between silent and active phases. Mapping these transitions offers a principled way to classify bursting dynamics and understand the mechanisms underlying pathological neural activity.

To generate bursting activity, the system must incorporate additional slow processes that modulate the excitability of the neuron over longer timescales. These slow variables influence the parameters of the fast subsystem, effectively controlling the timing and occurrence of bifurcations at the onset and offset of the bursts (Jirsa et al., 2014 [18]; Izhikevich, 2007 [17]).

The model used in this study captures these dynamics at the level of a single neuron immersed in an external potassium bath and is described by a system of four coupled differential equations (Depannemaecker et al., 2022 [7]). The fast subsystem consists of the membrane potential and a potassium gating variable, which govern rapid spiking behavior. The slow subsystem includes the intracellular potassium concentration and the extra-cellular potassium exchange with the surrounding bath. These slow variables evolve on a longer timescale and modulate the fast dynamics, thereby enabling the system to transition between silent and active phases via distinct bifurcation mechanisms.

Importantly, varying the extracellular potassium concentration alters the slow dynamics and can drive the fast subsystem through different onset–offset bifurcation combinations. These bifurcations are typically identified through analysis of the full system, requiring access to both fast variables—in this case, the membrane potential and the gating variable. However, in experimental settings, only the membrane potential is usually recorded. This motivates the need for methods that can infer bifurcation types using membrane potential data alone.

The present model reproduces a variety of bursting patterns observed in epilepsy, and these can be qualitatively classified based on established bifurcation signatures described in the literature (Jirsa et al., 2014 [18]; Izhikevich, 2007 [17]; Saggio et al., 2020 [31]). By comparing model outputs to known dynamical patterns (i.e. bifurcation features), it is possible to identify the likely bifurcation structures underlying seizure-like transitions. This approach enables classification of bursting regimes even in the absence of full access to all model variables, making it applicable to both computational and experimental data.

In this framework, different types of bifurcations can be identified based on specific features of the membrane potential trace. Saddle Node (SN) and Saddle Homoclinic (SH) bifurcations are distinguished from Saddle Node on Invariant Circle (SNIC) bifurcations at the onset or offset of bursting by the presence of a sudden shift in baseline potential or by measuring the difference between the pre-burst baseline and the recovery potential following a spike. In contrast, a Supercritical Hopf (SupH) bifurcation is identified by a gradual increase (or decrease) in spike amplitude during the onset (or offset) of the burst (Jirsa et al., 2014 [18]; Izhikevich, 2007 [17]; Saggio et al., 2020 [31]). By combining these onset and offset features, we are able to assign each time series to a specific bifurcation category.

For example, spike trains (Fig.2.c) show SNIC bifurcations at both onset and offset. In seizure-like events (SLE), multiple bifurcation combinations can be observed. The most common patterns are associated with two-dimensional bifurcation sequences such as SN–SH (Fig.2.b) or SN–SupH and SupH–SH (Fig.3.b). Both tonic spiking (TS) and sustained ictal activity (SIA) are characterized by continuous spiking without an offset phase (here referred to as “infinity offset”). These two patterns can be distinguished by their onset bifurcation: SNIC for TS (Fig.2.d), and either SN or SupH for SIA (Fig.3.c). Additionally, depolarization block can be identified by an initial SN–SupH burst, followed by a silent phase in which the membrane potential remains elevated above the baseline (Fig.3.d).

**Figure 1:**
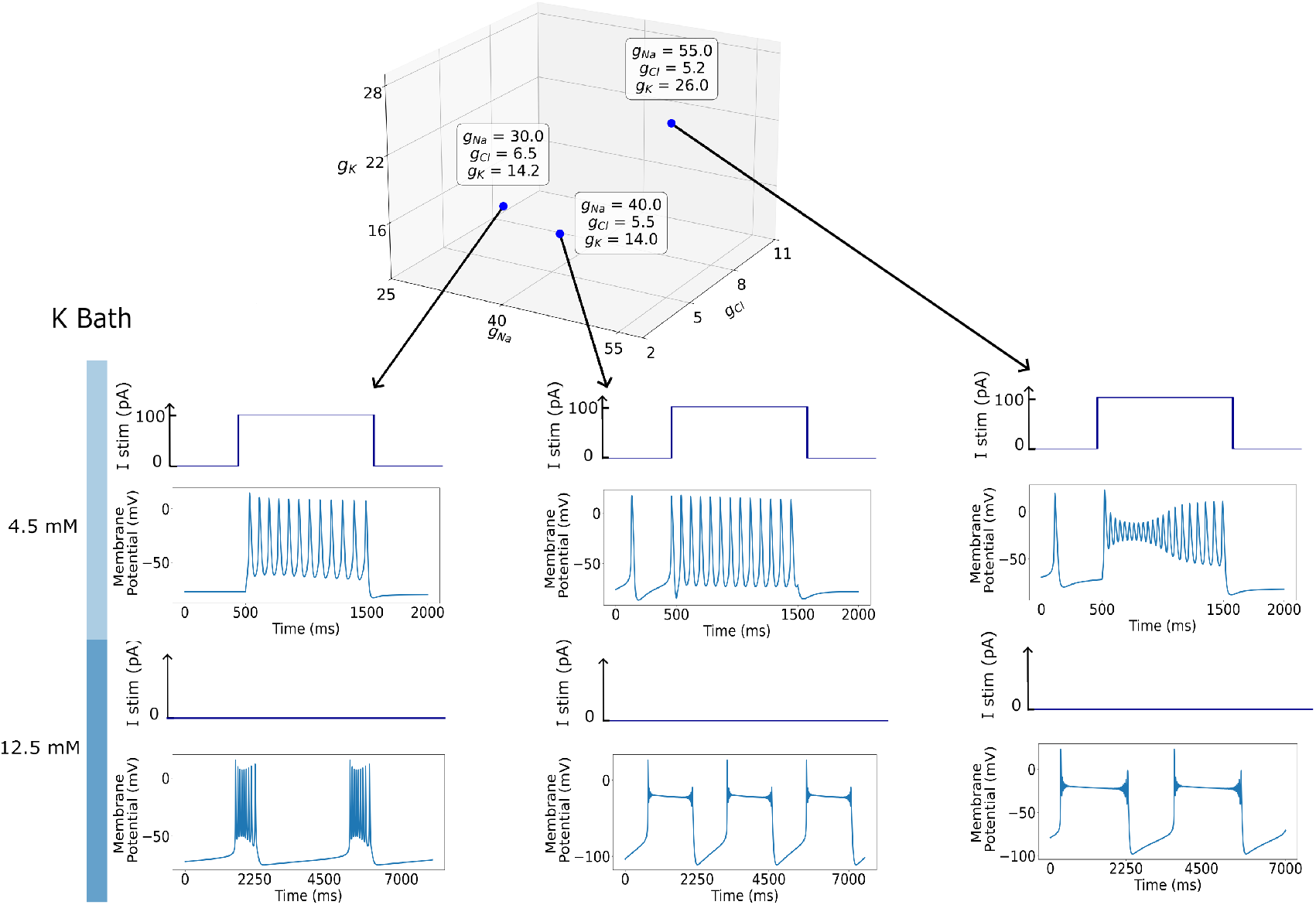
Membrane potential time series simulated under two different potassium bath concentrations and conductance parameter sets. Top panel: Three points are selected in the (*g*_Na_, *g*_Cl_, *g*_K_) parameter space. Middle row: At a potassium bath concentration of 4.5 mM, the neuron is in a resting state and is stimulated with a square-shaped injected current pulse of 100*pA*. The fist parameter sets produce similar membrane potential responses while the one on the right is different. Bottom row: At a higher potassium bath concentration of 12.5 mM, no external current is applied. The system exhibits spontaneous bursting activity, with qualitatively different dynamics: the left trace corresponds to a saddle-node/saddle-homoclinic (SN-SH) onset-offset bifurcation pattern, while the middle and the right trace correspond to a saddle-node/ Supercritical — Supercritical Hopf/saddle-homoclinic (SN-SupH—SupH-SH) bursting mechanism.

**Figure 2:**
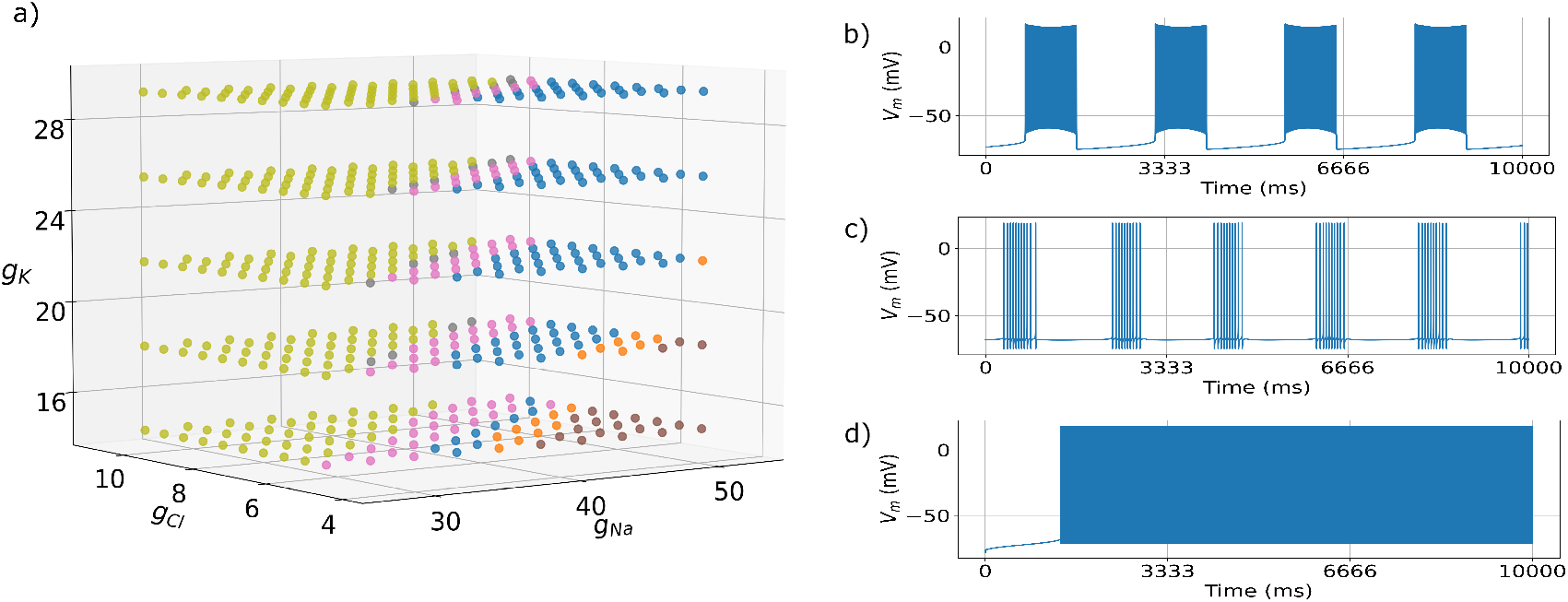
Simulation results for the external potassium bath fixed at 7.5 mM. (a) Three-dimensional plot in the space of the conductances at a fixed potassium level. Each point corresponds to a simulation of the model, colored according to the type of pattern observed in the time series. The categories identified at this potassium level are: SNIC-SNIC (grey); SNIC-Infinity (pink); SN-Infinity (blue); SN-SH (orange); and SN-SupH | SupH-SH (brown). Light green points correspond to simulations in which no signals emerge from the time series. Examples of membrane voltage time series corresponding to different patterns are shown in the right panels: (b) SN-SH; (c) SNIC-SNIC; (d) SNIC-Infinity.

**Figure 3:**
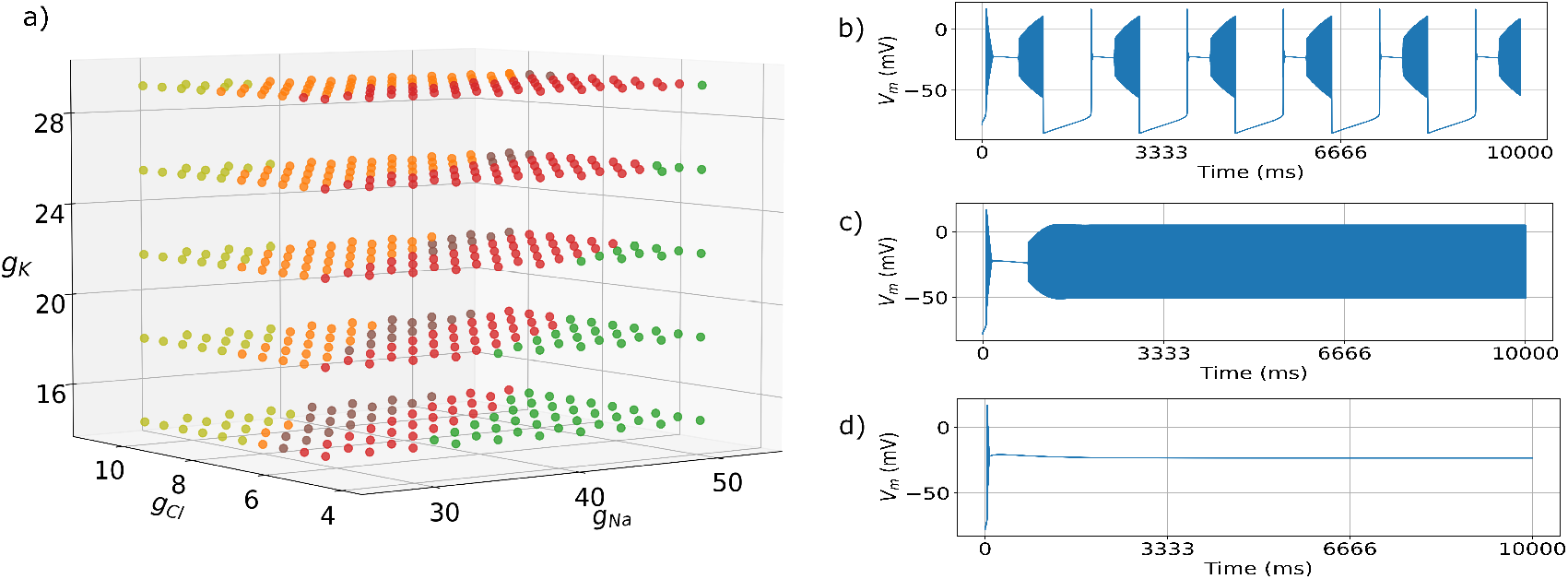
Simulation results for the external potassium bath fixed at 17.5 mM. (a) Three-dimensional plot in the space of the conductances at a fixed potassium level. Each point corresponds to a simulation of the model, colored according to the type of pattern observed in the time series. The categories identified at this potassium level are: SN-SupH|SupH-Infinity (red); SN-SupH (dark green); SN-SH (orange); and SN-SupH|SupH-SH (brown). Light green points correspond to simulations in which no signals emerge from the time series. Examples of membrane voltage time series corresponding to different patterns are shown in the right panels: (b) SN-SupH|SupH-SH; (c) SN-SupH |SupH-Infinity; (d) SN-SupH.

In this study, we first manually classified the model-generated time series according to their bifurcation structure. Based on this classification, we then developed a deterministic algorithm capable of qualitatively identifying bifurcation types using only the membrane potential (see Section 5). This allowed us to systematically explore the range of dynamical patterns generated by the single-neuron model as its biophysical parameters varied. In particular, we aimed to determine whether different parameter combinations could give rise to the same onset–offset bifurcation structure—that is, to explore degeneracy in the model’s dynamical behavior. These manually labeled features (see Materials and Methods) served as the foundation for developing an automated classification algorithm, which enabled the systematic analysis of bifurcation patterns across the full set of simulations.

We conducted an initial set of simulations comparing the output of two distinct parameter sets under varying conditions, including changes in extracellular potassium concentration and applied current. We then extended this analysis by systematically varying the three maximal conductances in the model, along with the extracellular potassium level (as summarized in Table1). This produced a dataset of 10,000 distinct model realizations, allowing us to study how degeneracy emerges and changes across different regions of parameter space.

**Table 1:** Parameter ranges and number of points for *g*_*Na*_, *g*_*Cl*_, *g*_*K*_, and [*K*]_*bath*_.

| Parameter | $g_{Na}$ | $g_{Cl}$ | $g_K$ | $[K]_{bath}$ |
| --- | --- | --- | --- | --- |
| Lower value | 26.67 | 5 | 14.7 | 7.5 |
| Upper value | 53.33 | 10 | 29.33 | 17.5 |
| Number of points | 10 | 10 | 5 | 20 |

### 3.2 Degenerate Parameters, Shared Patterns

Biophysical models of neural activity are defined by multiple parameters that represent key cellular properties. Among these, conductance parameters typically reflect average values derived from the many voltage-gated ion channels present in a single neuron (Hille, 2001 [15]). These ion channels can dynamically modify their electrophysiological characteristics through regulated synthesis and degradation, replacing existing channels with newly produced variants that differ in subunit composition, localization, or post-translational modifications (Marder and Goaillard, 2006 [22]). Such temporal fluctuations, combined with intrinsic variability across neurons, necessitate analyzing models over a broad range of conductance parameter combinations.

Variations in conductance values can significantly alter the complex mech-anisms underlying neuronal electrophysiological behavior (Alonso and Marder, 2019 [1]; Prinz et al., 2004 [28]; Foster et al., 1993 [11]). Different parameter sets, representing distinct internal neuronal configurations, may produce similar activity patterns under certain external conditions. However, when external conditions change, these internal differences can lead to qualitatively different neuronal responses (Fig. 1), although in some cases, comparable outputs emerge despite parameter variability, illustrating degeneracy.

Our analysis revealed that, even at a fixed extracellular potassium concentration, the model can exhibit a variety of activity patterns. This indicates that degeneracy observed under certain conditions can break down under others. Using the second set of simulations, we investigated how conductance parameter combinations that produce identical onset–offset bifurcations are distributed within the conductance space. For each fixed level of extracellular potassium ([*K*]_bath_), we examined the distribution of pattern types across conductance values.

At a given potassium concentration (Fig.2.a), up to six distinct activity patterns were observed. Parameter sets that produced the same pattern were found to be closely grouped, suggesting that these neuronal configurations share similar intrinsic properties. Within each pattern category, no further sub-clustering was detected, indicating the presence of compact degeneracy regions in conductance space—defined as connected subsets of parameters that give rise to the same pattern. We then examine how these degeneracy regions change across different levels of extracellular potassium. In this study, we distinguish between pattern classes and degeneracy regions, which describe different but related aspects of the system’s dynamics. A pattern class refers to a specific sequence of bifurcations—such as SNIC–SH or SN–SupH—that defines the qualitative features of neuronal activity under given external conditions. These classes are determined by the model parameters but do not uniquely identify them; in other words, different combinations of intrinsic conductances can give rise to the same pattern class. These combinations form a degeneracy region, which is the subset of parameter space where all points generate the same type of dynamical behavior.

While both concepts describe how neuronal activity is organized, their roles differ: pattern classes describe the type of dynamics expressed by the system, whereas degeneracy regions capture the structural variability—at the level of conductance parameters—that supports those dynamics. Distinguishing between these two helps clarify how different neuronal configurations can exhibit similar activity under specific conditions, and how their responses may diverge when external factors change.

Previous studies have shown that increasing extracellular potassium concentration in neurons with fixed conductances can lead to qualitative changes in activity patterns (Depannemaecker et al., 2022 [7]). Here, we investigate whether this effect is uniform across different combinations of conductance parameters.

At the highest potassium concentration tested, new activity patterns emerged that were not present at lower levels, while others disappeared (Fig.3.a). Although the degeneracy regions remained compact, their distribution across pattern categories differed from that observed at lower potassium levels (cf. Fig.2.a). This indicates that the organization of degeneracy regions is sensitive to changes in external conditions.

Overall, our results show that both intrinsic neuronal properties and external ionic concentrations jointly determine the activity patterns exhibited by the model. Increasing potassium modulates the number and identity of conductance configurations associated with each type of pattern. In particular, the proportion of patterns typically associated with pathological activity increases with higher potassium levels, while some patterns observed under baseline conditions are no longer reproducible at elevated concentrations. These shifts suggest that the structural similarity within degeneracy regions is condition-dependent: under varying external conditions, internal variability gives rise to different pattern distributions (Alonso and Marder, 2019 [1]; Alonso and Marder, 2020 [2]). In the biological system, this corresponds to changes in neuronal conductance profiles, while in the model, it is reflected by altered compositions of the degeneracy regions.

### 3.3 External Conditions Shape Neuronal Responses

Transitions between activity pattern categories do not occur uniformly across all neuronal configurations. While similar intrinsic properties may underlie pattern degeneracy under fixed external conditions, differences in these properties can result in divergent responses when the extracellular potassium concentration changes. To fully characterize each neuronal realization, it is necessary to examine its behavior across the entire range of potassium concentrations and assess how the expression of activity patterns varies with these changes.

Across simulations, model realizations exhibit different numbers of distinct activity patterns as the extracellular potassium level increases. Neurons that express the same number of patterns (Fig.4) do not necessarily form compact regions in conductance space, unlike the degeneracy regions observed at fixed potassium levels. The only clearly delineated group consists of parameter sets that fail to generate any activity across the full potassium range—these form a well-defined region in parameter space.

**Figure 4:**
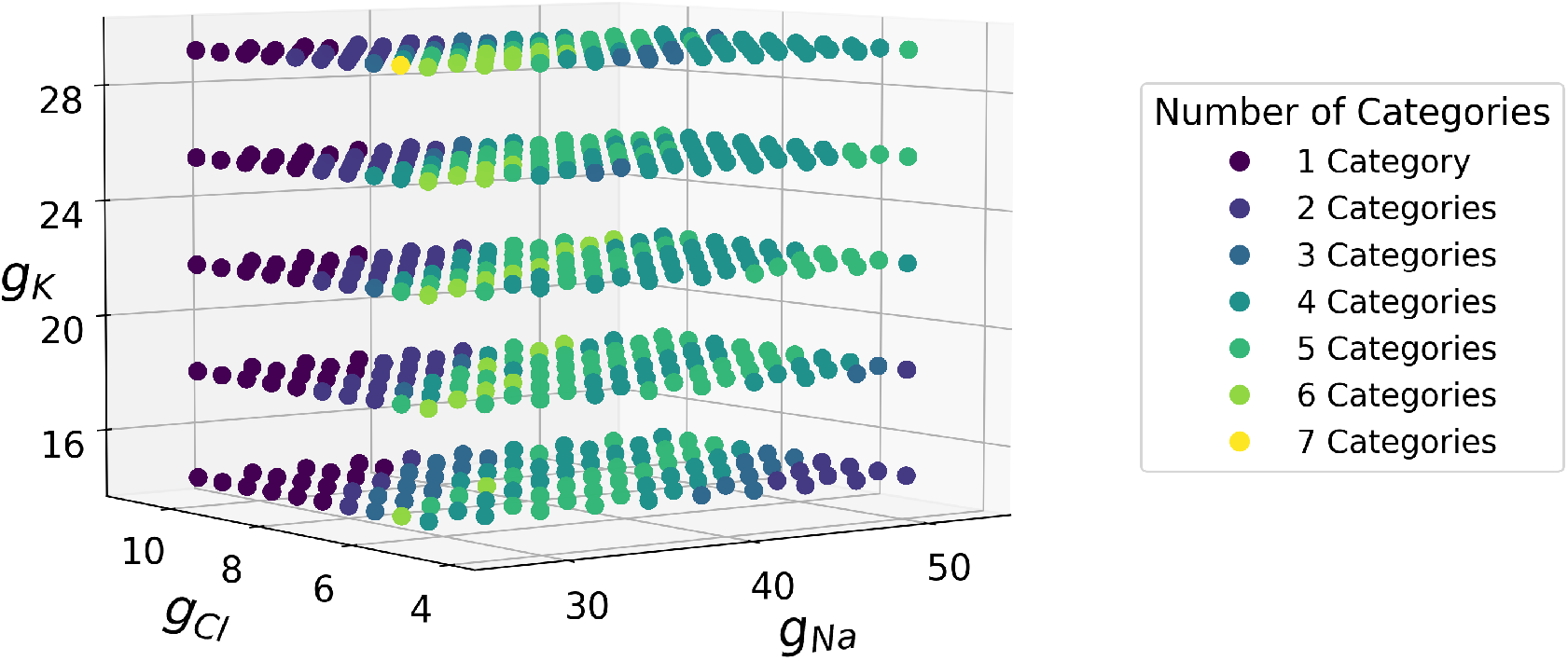
Representation of the number of crossing categories through the potassium bath in parameter space. Three-dimensional plot where each point represents a value in the conductance space, simulated for multiple potassium bath levels within a range from 7.5 to 17.5 mMol. The colors of the points indicate the number of distinct categories that each point can exhibit within the considered potassium range.

This finding implies that neurons exhibiting the same activity pattern under a given condition may diverge when external conditions change. Furthermore, neurons that follow the same sequence of pattern transitions (i.e., identical paths in pattern space) may still differ in the specific potassium concentrations at which these transitions occur. As shown in Fig.5, two parameter sets may display identical patterns up to a certain potassium level, after which they diverge and express different behaviors. For instance, the model realization corresponding to point 1 remains silent across all potassium levels. In contrast, points grouped as 2, 3, 4, 5 and 8, 9 exhibit identical sequences of patterns but over different potassium intervals. Other neurons, such as those labeled 6 and 7, initially express similar patterns at low potassium levels, but then diverge into different pattern categories as potassium increases.

**Figure 5:**
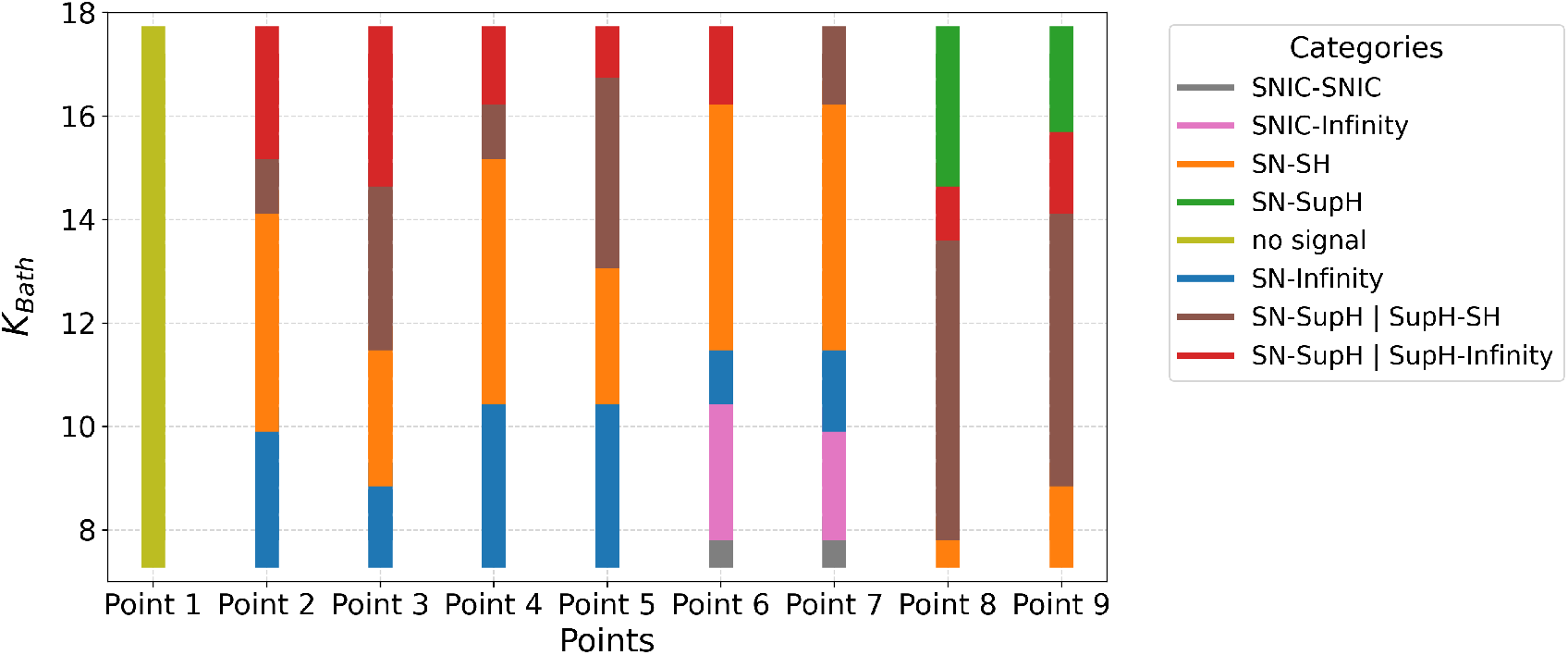
Transition paths among categories across the analyzed potassium range for 9 distinct points in the conductance space. In the colorbars, each color denotes a specific activity pattern, while the length of the colored segment represents the external concentration interval over which the neuron exhibits that pattern.

These results demonstrate that different conductance configurations can lead to distinct dynamical responses, even when they produce the same pattern under baseline conditions. More broadly, the transition between activity categories is not fixed and can vary with external potassium concentration. Even when the pattern trajectory remains the same, the specific potassium levels required to induce transitions between patterns may differ.

In summary, this variability suggests that each point in conductance space has a unique sensitivity profile to external perturbations. Therefore, to better understand the mechanisms underlying such differences, we next investigate which specific conductance features contribute to greater or lesser resilience—defined here as the ability of a given realization in parameter values, to resist changes in activity pattern as extracellular potassium concentration increases.

### 3.4 Resilience Reflects Ion Channel Configuration

Because of intrinsic differences, neurons may exhibit the same pattern class at different levels of extracellular potassium concentration ([*K*]_*bath*_). We define the resilience of a neuron as its resistance to changes in activity pattern when external conditions are modified. Variability in resilience reflects how changes in internal properties—such as ionic conductances—influence the neuron’s ability to maintain its current behavior. The goal of this analysis is to identify which intrinsic conductances most strongly determine this resilience.

In our model, we hypothesize that certain conductances, in particular potassium conductance, have a greater influence on resilience than others. Since the activation and termination of bursts are controlled by potassium fluxes between the external bath and the extracellular space, we expect that the potassium conductance plays a dominant role in determining how much [*K*]_*bath*_ must increase to induce a pattern transition.

To test this, we conducted the following analysis: for each point in conduc-tance space that expresses a specific pattern class, we recorded the minimum value of [*K*]_*bath*_ required to elicit that pattern. Then, we calculated the difference in this minimum value between each point and its nearest neighbors along each axis of the three-dimensional conductance space. For each non-boundary point, this yielded a vector whose components reflect the change in resilience with respect to each conductance direction. The orientation of these vectors indicates which conductances most influence resistance to external perturbations.

In both the Depolarization Block (Fig.6.a.1) and the SN–SH class of Seizure-Like Events (Fig.6.b.1), the vectors are predominantly aligned along the potassium conductance axis, supporting the idea that resilience increases most strongly with potassium conductance. Near the edges of the parameter space in Fig. 6.b.1, deviations from this trend are observed. These outliers likely result from classification inaccuracies, especially when bifurcation features become hard to distinguish near transitions between pattern classes.

**Figure 6:**
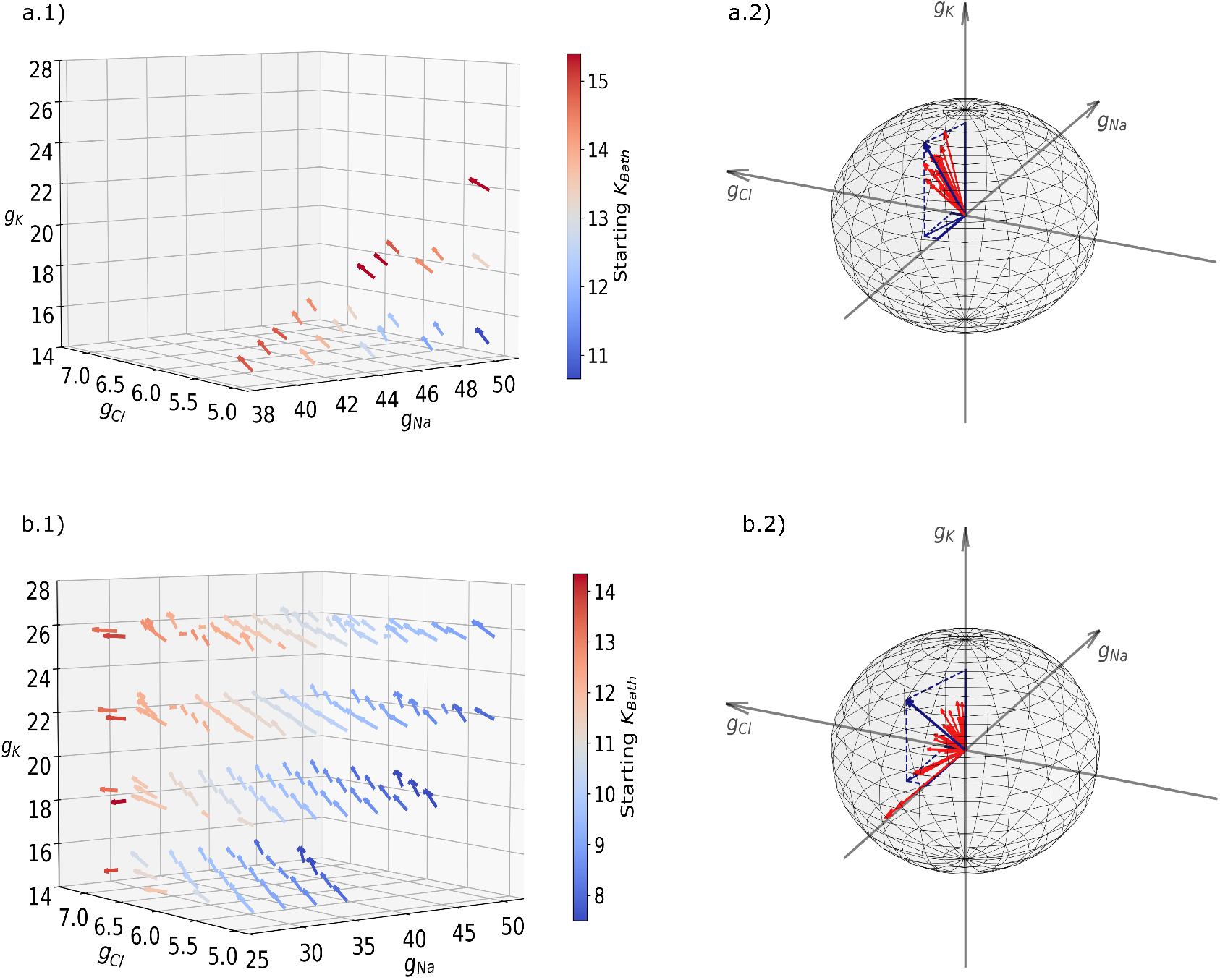
Vectors representing increasing resilience in the conductance space. Panels (a.1) and (b.1) present 3D plots for DB and SLEs (SN-SH), respectively. Each data point is depicted by an arrow whose color indicates the minimal potassium level required for inclusion in that category. Panels (a.2) and (b.2) display the corresponding vectors (red arrows) and their mean vectors (blue arrows) expressed in spherical coordinates.

From the spherical representation in Fig.6.a.2 and in Fig.6.b.2, we can observe that the vectors are clustered within a restricted portion of the sphere. The angular coordinates of the average vector are given by (*r* = 3, *θ* = 32^°^,*ϕ* = −168^°^) for Depolarization Block (SN-SupH), and (r=3, *θ* = 30^°^, *ϕ* = −159^°^) for SN-SH. Here, *θ* represents the polar angle measured from the *g*_*K*_ axis, and *ϕ* is the azimuthal angle measured in the *g*_*Na*_–*g*_*Cl*_ plane from the positive *g*_*Na*_ axis. According to spherical coordinates, the Cartesian components can be recovered using the inverse transformations:

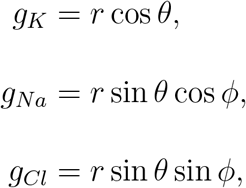

where *θ* ∈ [0, 2*π*] is the polar angle, *ϕ* ∈ [0, 2*π*] is the azimuthal angle, and *r* is total conductance magnitude. The analysis of the direction vectors confirms that the potassium conductance (*g*_*K*_) is the dominant contributor to resilience. The *g*_*K*_ component is consistently positive and larger than the others, indicating that increasing potassium conductance strengthens the neuron’s resistance to transitions in activity patterns. In contrast, the contributions of sodium (*g*_*Na*_) and chloride (*g*_*Cl*_) conductances are smaller and typically negative, suggesting that higher values of these conductances tend to reduce resilience. Among the two, *g*_*Cl*_ has the weakest influence, consistent with its limited role in this model.

These findings align with the model’s structure and assumptions. In conductance-based models, each ionic current depends on both the conductance value and the difference between the membrane potential and the ion’s Nernst potential. The Nernst potential is itself determined by the ratio of extracellular to intracellular concentrations. Importantly, in this model, potassium dynamics are modulated by both slow variables—one governing intracellular potassium accumulation and the other controlling uptake from the external bath [7]. Sodium concentration, in contrast, varies in opposition to potassium, and is affected by only one of the slow variables, while chloride concentration is held constant.

As a result, increasing potassium conductance effectively compensates for external perturbations by amplifying the current associated with the main regulatory pathway. For sodium, the compensatory effect arises from decreasing conductance, due to the inverse relationship between sodium and potassium concentrations. The small influence of chloride conductance stems from its static concentration and the fact that its effect on membrane potential depends on the sign of its charge and the overall potential difference.

Altogether, this analysis highlights how the interplay of ionic concentrations and conductance values shapes the neuron’s electrophysiological behavior. Despite this complexity, multiple distinct configurations of conductances can give rise to similar activity patterns, particularly those governed by onset–offset bifurcations.

## 4 Discussion

In this study, we developed a method to classify neuronal time series based on features of their bursting dynamics, with the goal of identifying characteristic onset–offset bifurcation types. These features correspond to bifurcations typically found in low-dimensional dynamical systems (Jirsa et al., 2014 [18]; Izhikevich, 2007 [17]; Saggio et al., 2020 [31]). Unlike classical bifurcation detection techniques—which require full knowledge of all system variables (Dhooge et al., 2003 [8]; Gast et al., 2019 [14])—our method relies solely on the membrane potential, making it applicable to experimentally recorded data. An important aspect of this work is that both pattern classification and the identification of degeneracy regions are performed using only the membrane potential (*V*_*m*_), which is typically the only variable accessible in experimental recordings. Unlike in computational models, where the full set of internal variables and parameters (e.g., ionic conductances, gating variables) is known, experimental data rarely provide direct access to the underlying biophysical configuration. As a result, the pattern class in this study is defined operationally based on features extracted from the membrane potential—such as spike amplitude evolution and baseline shifts—rather than from direct bifurcation analysis involving all system variables. Likewise, degeneracy regions here refer to subsets of parameter space inferred to produce the same voltage-based pattern, even though the full dynamical system structure remains hidden in real biological neurons. This distinction reinforces the relevance of the proposed method for experimental contexts: it provides a framework to classify activity patterns and explore their parametric origins using only observable voltage traces, thereby offering a bridge between theoretical models and electrophysiological data.

We applied this method to a single-compartment neuron model and sys-tematically varied its intrinsic properties by tuning three maximal ionic conductances (sodium, potassium, and chloride), while holding the external potassium concentration fixed. We found that different combinations of conductances could give rise to the same activity pattern and bifurcation sequence. This observation is a manifestation of degeneracy: the capacity of structurally distinct configurations to produce functionally equivalent outputs (Edelman and Gally, 2001 [9]).

Degeneracy has been widely observed in model of the nervous system, particularly in conductance-based models (Alonso and Marder, 2019 [1]; Alonso and Marder, 2020 [2]; Migliore et al., 2018 [23]). Our results extend this principle to the level of dynamical transitions associated with pathological states. Specifically, we show that a given epileptiform activity pattern—such as bursting—can emerge from multiple, topologically compact regions in conductance space. Within each region, certain conductances are tightly constrained to support the observed dynamics, while others exhibit wider variability, consistent with previous reports of parametric compensation in ionic currents (Migliore et al., 2018 [23]). To investigate how these degenerate configurations respond to changes in the external environment, we then varied the extracellular potassium concentration. This manipulation mimics a well-known pathological driver of excitability shifts in epilepsy and other disorders. While degeneracy was preserved under this perturbation, the internal conductance profiles supporting each pattern were redistributed. In particular, the conductance values that led to similar dynamics at baseline often diverged in their reactivity to increasing potassium levels.

To quantify this divergence, we introduced the notion of resilience, defined as the minimum extracellular potassium concentration required to elicit a given activity pattern from a particular conductance configuration. This metric captures the threshold for transition into a specified regime (e.g., from tonic firing to bursting) under external drive. Importantly, it allows us to evaluate not just whether a configuration supports a pattern, but how sensitive it is to perturbations in the extracellular environment. Using a directional analysis across the three conductance axes, we estimated local gradients of resilience and visualized these changes using a spherical vector representation. Our findings show that potassium conductance plays the largest role in shaping resilience, while sodium and chloride have smaller and more asymmetric effects—especially at low conductance values.

These results reveal that degeneracy is not a static property: even if multiple conductance configurations are equivalent under fixed conditions, their trajectories under external modulation may diverge. Thus, degeneracy co-exists with structured variability in how neurons respond to environmental change. This has important implications for both modeling and interpretation of electrophysiological data.

Our approach, while targeted and informative, involves a number of simplifications. We varied only three maximal conductances and kept all other parameters—including those controlling interactions with the extracellular space—constant. The conductances used represent averaged properties of ion channels rather than detailed channel kinetics, limiting the biological variability captured. Moreover, our model includes only potassium as the extracellular modulator, excluding other ion concentrations, synaptic input, or metabolic effects. These constraints narrow the range of observed mechanisms and the possible manifestations of resilience.

In larger models with more dimensions, the degeneracy landscape becomes increasingly complex and difficult to visualize. In such cases, methods from machine learning—especially Bayesian inference—can support structured exploration of parameter space, quantify uncertainty, and identify high-dimensional degeneracy manifolds (Baldy et al., 2025 [4]). We have proposed a framework to identify and analyze degeneracy in neuronal activity patterns, particularly those relevant to epileptic dynamics. Our method enables the classification of bursting regimes using membrane potential alone and introduces a metric—resilience—to quantify how internal configurations modulate a neuron’s response to extracellular perturbations.

By varying conductances in a simplified neuron model, we confirmed the presence of degeneracy across multiple bifurcation types under fixed external conditions. We further demonstrated that while degenerate configurations support similar patterns at baseline, they differ in how they transition under changes in potassium concentration. This dual analysis of pattern robustness and sensitivity provides a new way to study intrinsic variability in neurons.

These findings have broader implications for experimental neuroscience. Electrophysiological recordings often reveal functionally similar neurons with diverse molecular or biophysical profiles. Our results suggest that comparing neurons solely based on molecular markers is insufficient (Northcutt et al., 2019 [25]); instead, their responses to perturbations offer a richer basis for classification (Kim et al., 2020 [19]). This is also relevant at the network level, where intrinsic conductance variability has been shown to affect dynamics more than synaptic weights.

Ultimately, a deeper understanding of degeneracy and resilience at the single-neuron level could inform personalized models of brain function and dysfunction. In epilepsy, in particular, variability in single-neuron resilience may contribute to heterogeneous responses to treatment and drug resistance (Price and Friston, 2002 [27]). Our approach provides a principled basis for such investigations and a step toward the integrative modeling of neuronal variability across scales.

## 5 Materials and Methods

### 5.1 Model

In this study, we adopt a single level neuron model that describes the electro-physiological dynamics of a neuron immersed in an extracellular potassium bath. The model is formulated as a system of four coupled ordinary differential equations forming a slow-fast dynamical system.

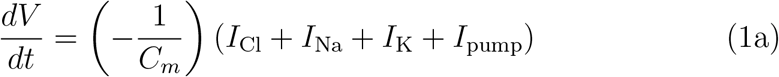

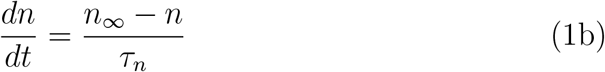

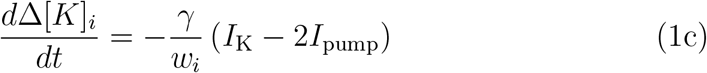

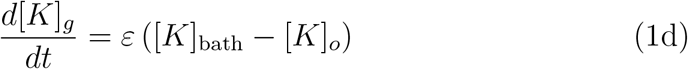

Eq. (1a) and Eq. (1b), describing the membrane potential and the gating variable for potassium conductance, respectively, constitute the fast subsystem. In contrast, the slow subsystem is defined by Eq. (1c) and Eq. (1d), which describe the dynamics of intracellular potassium concentration and extracellular potassium buffering by the external bath, respectively. The ionic currents are given by the following expression:

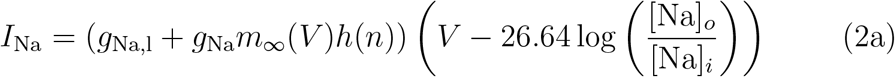

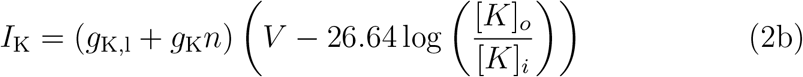

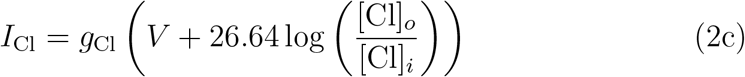

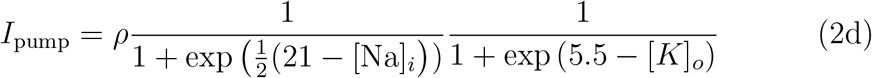

Eq. (2a), Eq.(2b), and Eq. (2c) describe the ionic currents flowing through the voltage-gated sodium, potassium, and chloride channels, respectively.

Eq. (2d) represents the current generated by the sodium-potassium pump. The conductance terms in these equations depend on the membrane potential *V* and the gating variable *n*, and are defined in Eq. (3a),Eq. (3b) and Eq. (3c):

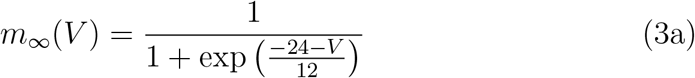

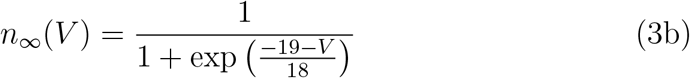

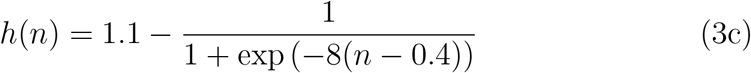

The initial values for the dynamical variables and the ionic concentration are shown in Tab. 2.

**Table 2:** Initial values of the model variables.

| Variable | Initial Value |
| --- | --- |
| $V$ | -70 mV |
| $n$ | $n_{\infty}(-70)$ |
| $\Delta[K]_i$ | 0 |
| $[K]_g$ | 0 |
| $[K]_o$ | $[K]_{0,o}$ |
| $[K]_i$ | $[K]_{0,i}$ |
| $[Na]_i$ | $[Na]_{0,i}$ |
| $[Na]_o$ | $[Na]_{0,o}$ |
| $[Cl]_i$ | $[Cl]_{0,i}$ |
| $[Cl]_o$ | $[Cl]_{0,o}$ |

The initial conditions for the ionic concentrations were chosen to reflect physiologically plausible values commonly used in computational models of neuronal dynamics. Specifically, the initial intracellular sodium concentra-tion was set to [Na]_*i*,0_ = 16.0 mM, while the extracellular sodium concentration was [Na]_*o*,0_ = 138.0 mM. For potassium, the intracellular and extracellular concentrations were initialized to [*K*]_*i*,0_ = 140.0 mM and [*K*]_*o*,0_ = 4.80 mM, respectively. Chloride concentrations were also included, with an extra-cellular value of [Cl]_*o*,0_ = 112.0 mM and an intracellular value of [Cl]_*i*,0_ = 5.0 mM. At each timestep, the ionic concentrations vary as follow:

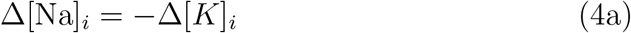

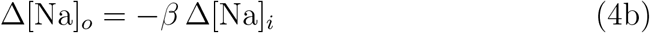

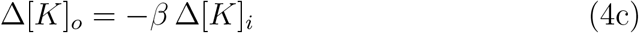

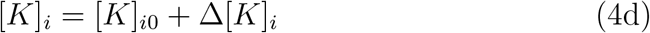

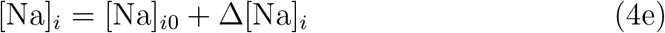

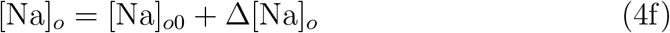

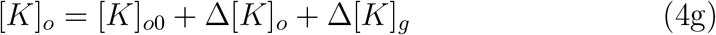

The conducatance parameters and the potassium concentration were varied in the intervals shown in Tab.1, while the other parameters are fixed at values in Tab.3.

**Table 3:** Model parameters fixed.

| Parameter | Symbol | Value |
| --- | --- | --- |
| Conversion factor | $\gamma$ | $0.04 \text{ mol}/(\text{C} \cdot \mu\text{m}^3)$ |
| Diffusion time constant | $\varepsilon$ | $0.01 \text{ ms}^{-1}$ |
| Membrane capacitance | $C_m$ | 1 nF |
| Gating time constant | $\tau_n$ | 0.25 |
| Intracellular volume | $w_i$ | $2160 \mu\text{m}^3$ |
| Extracellular volume | $w_o$ | $720 \mu\text{m}^3$ |
| Intra/extra cellular volume ratio | $\beta$ | 3 |
| Maximal Na/K pump current | $\rho$ | 250 pA |
| Sodium leak conductance | $g_{\text{Na},l}$ | 0.02 nS |
| Potassium leak conductance | $g_{\text{K},l}$ | 0.12 nS |

The simulations themselves are based on the integration of the system of differential equations using the Runge-Kutta method. They are performed under fixed initial conditions and span a time range of 10,000 ms. This approach ensures that the dynamic behavior is accurately captured for each set of parameters.

### 5.2 Manual targeting procedure of onset-offset bifurcations

In this work, we first developed a Python tool to classify bifurcations by visually estimating the main features related to onset-offset bursts (Izhikevich 2007 [17]; Jirsa et al. 2014 [18]; Saggio et al. 2020 [31]). According to the model (Depannemaecker et al. 2022 [7]), only one pattern (dynamotype) appears in a given time series. For an accurate estimation, we required time series with at least three spikes. We did not consider the first burst because its onset-offset bifurcations can be influenced by the initial conditions of the system.

The manual targeting tool works as follows. It starts from an initial point in a three-dimensional parameter space defined by the three conductances, while keeping the potassium concentration fixed. The tool displays the membrane potential plots on the screen. Next, the user presses the button corresponding to the observed dynamotype. Once a button is pressed, the current parameter values and the selected target are recorded in a data file. After saving the data, the program automatically runs the simulation for the next set of conductance values.

This procedure was repeated for the entire range of conductance values and was applied for each potassium concentration listed in Tab.1.

### 5.3 Deterministic classifier algorithm

After manually classifying the time series according to the observed patterns, we collected the most relevant features to develop a deterministic algorithm for classifying the time series. First, we analyzed the time series based on the spikes they emitted. For spike detection, we used the SciPy find peaks function. Initially, we identified all time series with no spiking activity and labeled them as “no signal” time series. For the remaining time series, we grouped spikes into bursts using an algorithm based on the inter-spike distance. For each time series, we computed—when available—the last two bursts. This approach helps us avoid the transient and less accurate behav-ior often observed in the first burst. For both bursts, we computed the onset and offset features. The full list of features is available on GitHub; here, we only mention those used for bifurcation detection.

Next, we divided the ensemble of time series into two main groups based on the presence or absence of the SupH bifurcation. According to the literature, spike amplitude shows an increasing trend at onset and a decreasing trend at offset. Therefore, we calculated the ratio between the spike amplitudes at onset (or offset) and the maximum spike amplitude in the burst. If the amplitude ratio at onset for the last burst is below 0.8, we classify the time series as exhibiting a SupH bifurcation. In cases where the last burst begins just before the end of the time series, we refer to the amplitude ratio of the penultimate burst.

Once we defined the SupH and non-SupH groups, we further distinguished the different pattern categories within each group. In the SupH group, if the number of spikes in the last burst exceeds 5000, the classification is SN-SupH |SupH-Infinity. For the remaining time series, if the burst is identified as type 1 (or type 2 combined with a variance in the peak amplitude of the last burst less than 20), we classify it as SN-SupH (Depolarization Block). All other time series in this group are classified as SN-SupH |SupH-SH.

For the non-SupH group, we used the following procedure. To distinguish time series with not ending spiking activity (Infinity offset) from those without, we computed the ratio of the number of spikes in the penultimate burst to the number in the last burst. If this ratio is less than 1, the time series is classified as Infinity offset. To further differentiate the onset bifurcation in both the Infinity and non-Infinity cases, we combined two features. First, we estimated the baseline jump, defined as the difference between the membrane potential measured just before the start of the burst and its minimum value within the interval defined by the first two spikes. Then, defining the spike period as the time difference between consecutive spikes, we computed the ratio between the first two spike periods to complement the baseline jump. If the baseline jump is less than 0 and the period ratio is greater than 1, the classification is SNIC-SNIC; otherwise, we label it as SN-SH.

The use of these features is consistent with the bifurcation characteristics reported in the literature. In fact, a baseline jump occurs at onset (or offset) only for the SN (or SH) bifurcation, and for the SNIC bifurcation, the frequency approaches zero at the bifurcation point, meaning that its inverse (the spike period) must be greater than 1.

### 5.4 Computation and Visualization of resilience Variations

To assess the resilience of specific points within the parameter space concerning the emergence of a particular pattern as the potassium bath concentration increases, we employed the following methodology. Initially, for each point in the parameter space, we determined the minimal concentration of potassium bath required to elicit the specified pattern. This threshold represents the critical potassium level at which the pattern first appears. Subsequently, for each point, excluding those situated at the boundaries of the selected region, we computed the differences in potassium bath concentrations between the point and its nearest neighbors along each axis of the three-dimensional conductance space. This process yielded a vector for each point with components that corresponded to these differences. The orientation and magnitude of this vector provide insight into the sensitivity of the pattern’s manifestation to variations in conductance parameters. Specifically, the vector’s direction indicates which conductance parameters most significantly influence the pattern’s resilience, while its magnitude reflects the degree of sensitivity, with larger magnitudes suggesting greater sensitivity to parameter changes.

To further analyze the orientation of these vectors within the conductance space, we converted them from Cartesian coordinates to spherical polar co-ordinates. In this coordinate system, each vector is characterized by three parameters: the radial distance *r*, the polar angle *θ*, and the azimuthal angle *ϕ*. The radial distance *r*, representing the vector’s magnitude, was calculated as

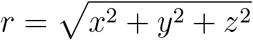

where *x, y*, and *z* are the vector’s components along the respective conductance axes. The polar angle *θ*, denoting the angle between the vector and the positive *z*-axis, was determined using

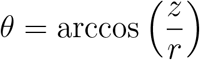

The azimuthal angle *ϕ*, indicating the angle between the projection of the vector onto the *xy*-plane and the positive *x*-axis, was obtained via

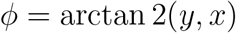

This transformation facilitated a more intuitive understanding of the vectors’ orientations, thereby elucidating the conductance parameters that most critically affect the resilience of the observed pattern.

## 6 Acknowledgments

The project leading to this publication has received funding from the Excellence Initiative of Aix-Marseille Université - A*Midex, a French “Investissements d’Avenir programme” AMX-21-IET-017. This research has received funding from the European Union’s Horizon Europe Programme under the Specific Grant Agreement No. 101147319 (EBRAINS 2.0 Project). It has also received funding from the European Union’s Horizon Europe Programme under the Specific Grant Agreement No. 101137289 (Virtual Brain Twin Project). This work has benefited from a government grant managed by the Agence Nationale de la Recherche (ANR) under the France 2030 program, reference ANR-22-PESN-0012.

## Information Sharing Statement

All code is available on GitHub (https://github.com/GianmarCafaro/Pattern_Detection/).

